# *vassi* – verifiable, automated scoring of social interactions in animal groups

**DOI:** 10.1101/2025.07.15.664909

**Authors:** Paul Nührenberg, Aneesh P. H. Bose, Alex Jordan

## Abstract

Behavioral biologists, from neuroscientists to ethologists, rely on observation and scoring of behavior. In the past decade, numerous methods have emerged to automate this scoring through machine learning approaches. Yet, these methods are typically specified towards laboratory settings with only two animals, or employed in cases with well-separated behavioral categories. Here, we introduce the *vassi* Python package, focusing on supervised classification of directed social interactions and cases in which continuous variation in behavior means categories are less distinct. Our package is broadly applicable across species and social settings, including single individuals, pairs and groups, and implements a validation tool to separate behavioral edge cases. *vassi* has comparable performance to existing approaches on a behavioral classification benchmark, the CALMS21 mouse resident-intruder dataset, and we demonstrate its applicability on a novel, more naturalistic and complex dataset of cichlid fish groups. Our approach highlights future challenges in extending supervised behavioral classification to more naturalistic settings, and offers a methodological framework to overcome these challenges.

**Lay Summary:** *vassi* (*verifiable, automated scoring of social interactions*) is a flexible, Python-based framework for automated behavioral classification and its verification through interactive visualization. *vassi* enables researchers to quantify directed social interactions in animal groups in naturalistic settings, bridging the gap between traditional ethology and modern computational tools.

## Introduction

The study of animal behavior has wide-ranging impacts across fields including conservation, animal welfare, neuroethology, genetics, and evolutionary biology. At the intersection of these fields lies the discipline of ethology, which at its core is concerned with the phenomenological, causal, ontogenetic, and evolutionary aspects of behavior in wild, freely moving animals (Tinbergen, 1963). Traditionally, ethologists have recorded behavioral events and intervals during experiments or observations using pen, paper, and a stopwatch (Altmann, 1974; Barton & Johnson, 1990), an approach that can be error-prone for various reasons. Observing multiple animals simultaneously requires humans to multitask, which is typically associated with performance costs (Poljac et al., 2018). The collection of temporal measurements of behavior during live observations is also likely affected by the speed-accuracy tradeoff, a well-studied phenomenon in humans and across taxa (e.g., reviewed in Heitz, 2014). Jeanne Altmann (1974) highlighted the challenges of manual behavioral sampling and scoring, sharing an anecdote that emphasized these difficulties: *“[…] with two observers, one 15-minute sample per hour was near the upper limit of our capacity when obtaining an accurate record, with some 5 dozen social behavior categories, of who did what to whom and in what order, as well as keeping track of most nonsocial behavior, durations, and time-out periods.”* Even when human observers are able to keep track of complex animal interactions, there can be considerable variation among observers, particularly with respect to start and stop times (Segalin et al., 2021). Moreover, animal behaviors may vary at subtle levels that are difficult for observers to reliably recognize or measure – they may occur at very short or very long time-scales (Berman, 2018), exist in sensory modalities outside of human scope (e.g., olfaction; Nielsen et al., 2015), or comprise signals that are too complex for an observer to interpret directly (Patricelli & Hebets, 2016). While some of these difficulties can be overcome in the laboratory through careful experimental design, observation of animals in more natural conditions, for example in larger groups, in complex environments, or indeed under natural conditions, can compound these issues.

In response to these challenges the sub-field of computational ethology has emerged, in which researchers employ data-driven methods to measure and analyze the behavior of animals (outlined in Anderson & Perona, 2014). Powered by the rapid expansion of video recording as a methodology, supervised machine-learning algorithms can be trained on trajectory, posture, and video data of animals to automatically detect and classify behaviors of interest (implemented in tools like JAABA, SimBA, MARS or A-SOiD; Kabra et al., 2013; Goodwin et al., 2024; Segalin et al., 2021; Tillmann et al., 2024), leveraging data from recent methods for single and multi-animal detection, tracking and posture estimation that achieve unprecedented detail in raw measurements of animal movement (e.g., DeepLabCut, DeepPoseKit, idtracker.ai or sleap; Mathis et al., 2018; Graving et al., 2019; Romero-Ferrero et al., 2019; Pereira et al., 2022). In brief, a workflow to automate behavioral scoring may consist of the following steps: Keypoints that mark an animal’s body parts can be precisely tracked throughout a video, describing the animal’s posture over time. This data can then be used to compute a high-dimensional description of the animal’s behavior (i.e., spatiotemporal and postural features; implemented for example in JAABA or SimBA). Additional features can also be computed in relation to other tracked animals to capture key aspects of social behavior, such as relative body poses, distances and angles. Subsequently, a machine-learning model (e.g., a decision tree classifier or an artificial neural network) may be fitted to this high-dimensional data, with known behavioral labels as the target variable (supervised learning, see e.g., Anderson & Perona, 2014). Such models trained on ground-truth behavioral data can then be applied to classify novel tracking-derived data, typically from members of the same species and in the same study contexts. Hence, researchers only need to manually score a small, but representative subsample of their entire video data, and can apply a fitted model on the remaining, potentially much larger dataset. This workflow eliminates some of the problems that are associated with manual scoring, such as errors due to scorer fatigue, or scorer bias and disagreement in the case of multiple scorers, while achieving or even exceeding the scoring accuracy of expert annotators (e.g., Segalin et al., 2021).

Nevertheless, these approaches face their own challenges (outlined in Kennedy, 2022), many of which are exacerbated as datasets become more complex with multiple animals in naturalistic settings. To facilitate the scoring, analysis and communication of animal behavior, researchers often impose distinct categories on potentially continuous behavior, which can result in disagreement as to when a behavior starts and stops, or when one behavioral category transitions to another (Gomez-Marin et al., 2014; Segalin et al., 2021). The natural variation in timescales over which behavior occurs can lead to inconsistent scoring of behavioral states, both within and among research groups, which in turn can lead to inconsistent classification by supervised models trained on these data (see Siegford et al. (2023) for a review on factors that impede effective automated behavioral analyses). These problems do not necessarily imply a drawback of the computational methodology, but rather highlight characteristics of the data – a behavioral state may correspond to a specific function (or ‘meaning’; Rendall et al., 2009) despite its underlying continuous variation, and the same or divergent forms (or postures; Berman, 2018) may have different effects on conspecifics depending on context or history. In African cichlid fish for example, a lateral posture can be used among rival males as part a threat display, or between males and females as part of a courtship display, and shares an underlying genetic architecture (Alward et al., 2020). As such, behavioral categorization may not only depend on the kinematic structure, but also on the identity of the participants themselves. Automated methods may therefore need to resolve the identity and directionality of social behaviors between particular actors and recipients, and depending on the context, may need to account for interactions among many individuals within a group. Perhaps because of these potential difficulties in generating robust behavioral categories, combined with a strong focus on model species, many available approaches are highly specified towards certain conditions (commonly two interacting animals in simple experimental arenas), which may complicate the generalization of such approaches and their implementation across contexts (Anderson & Perona, 2014; Goodwin et al., 2020; Kennedy, 2022).

As the call for more naturalistic experiments becomes stronger (Bordes et al., 2023; Smith & Pinter-Wollman, 2021), so too does the need for adequate quantitative tools. Here, we present the *vassi* Python package (*verifiable, automated scoring of social interactions*), enabling researchers to perform behavioral classification outside the boundaries of the existing tool-landscape. With continuous advances in animal tracking tools, such as their expansion to naturalistic settings (with aerial drones or SCUBA diving; Koger et al., 2023; Francisco et al., 2020), researchers can generate more complex tracking datasets by increasing variables such as the number of tracked animals or the diversity of behavioral interactions. With our package, we provide software that is applicable to such diverse use-cases; *vassi* does not impose constraints on the number of individuals that may socially interact in a group, and allows measurement of the direction of behavioral interaction. The package also allows users to set or automatically tune the confidence threshold for behavioral categories, and to review the resulting behavioral predictions to verify their correctness. The latter aspect becomes essential in cases where behavioral categories themselves are not easily distinguishable and require further downstream verification of potentially important edge-cases. We validated our package on an existing benchmark dataset, the CALMS21 dataset of dyadic mouse interactions, demonstrating classification performance on par with other, more specialized software published alongside the benchmark (Sun et al., 2021). We further tested the package on a novel dataset of larger groups of freely interacting fish, showcasing its applicability in more naturalistic and complex scenarios with subtle behavioral variation.

## Results

### CALMS21 dataset

We validated the *vassi* package on an established benchmark for behavioral classification, the CALMS21 dataset (Sun et al., 2021). This dataset consists of individually tracked dyadic resident-intruder interactions in mice and corresponding behavioral annotations that assign each video frame to one of four categories (‘attack’, ‘investigate’, ‘mount’, or ‘other’). We trained an XGBoost classifier (Chen & Guestrin, 2016) to distinguish between these categories using the data processing and sampling pipeline implemented by our package (see methods section for details, XGBoost classifier with default parameters, number of boosting rounds set to 1000). We used this classification model for predictions on the CALMS21 test dataset (Figure 1). Then, we evaluated the model on this dataset by computing macro F1 scores, a commonly used performance metric for classification on imbalanced datasets that is calculated as the harmonic mean of precision and recall for each category, and then averaged across categories (Fawcett, 2006). Our base model achieved a per-frame macro F1 score of 0.804 ± 0.001 (mean and standard deviation of 20 pipeline runs with different random states) on the three foreground categories (‘attack’, ‘investigate’ and ‘mount’), and 0.840 ± 0.001 on all four categories (Figure 1C and D).

**Figure 1:**
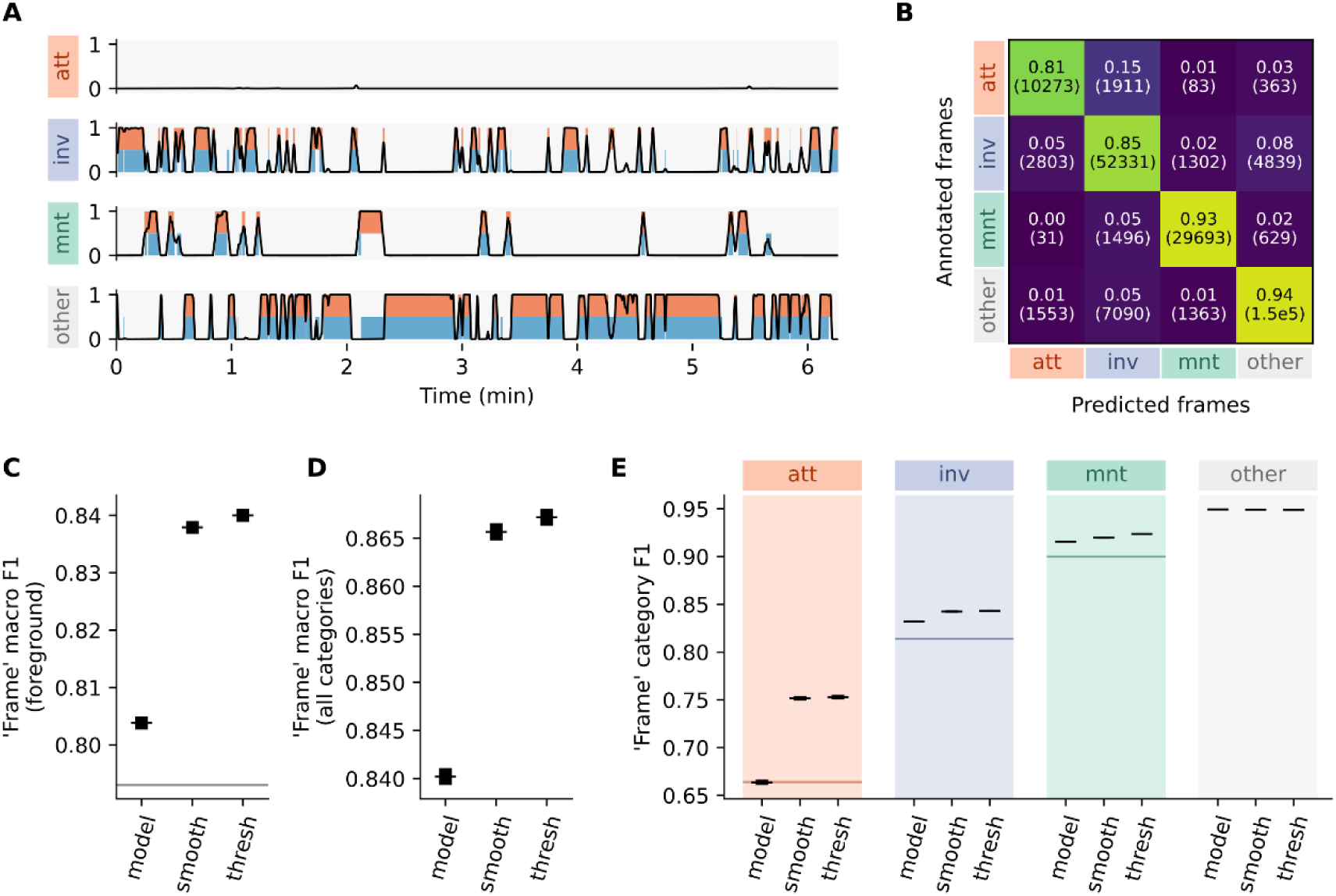
CALMS21 test dataset results. Behavioral categories are abbreviated: ‘attack’ – ‘att’, ‘investigation’ – ‘inv’ and ‘mount’ – ‘mnt’. All values represent means of 20 pipeline runs with different random states, standard deviations are shown as error bars if applicable. **A:** Behavioral timeline with ground-truth intervals (annotations, lower bars, blue) and predictions (upper bars, orange) for the four behavioral categories. Lines represent per-category model outputs (classification probabilities after smoothing). The last resident-intruder sequence of the test dataset is visualized, see SI Figure 5 for all 19 test dataset sequences. **B:** Per-frame confusion matrix of the four behavioral categories. Within each cell, upper values show the proportion of frames in agreement with annotated, ground-truth data (normalized across each row), lower values show the absolute frame counts. Results were visualized after model output smoothing and thresholding. **C and D:** Per-frame unweighted average (macro) F1 scores of raw model outputs (‘model’), after smoothing (‘smooth’), and after thresholding (‘thresh’), across only the behavioral foreground categories and across all four categories. **E:** Per-frame F1 scores for each category, calculated on raw model outputs, after smoothing, and after thresholding. **C and E:** Horizontal lines mark the F1 scores of the baseline model as reported in Sun et al. (2021).

We then transformed the raw model outputs (i.e., classification probabilities for each category and video frame) into accessible data in tabular form as behavioral events between a start and stop timestamp. We used two post-processing steps implemented by our package for this data transformation, temporal smoothing of model outputs and thresholding (see methods section for details). This post-processing entailed two sets of so-called hyperparameters that can affect predictive performance: Firstly, the kind (e.g., mean or median) and size of a temporal smoothing window when applying a filter to model outputs, and secondly, category-specific decision thresholds. We conducted hyperparameter optimization to find optimal values for these additional parameters. Smoothing model outputs using these values improved the quality of classifications (per-frame F1 score of 0.838 ± 0.001 on foreground categories, 0.866 ± 0.001 on all categories), outperforming the primary results without smoothing (Figure 1C and D). Thresholding these smoothed results with category-specific decision thresholds yielded an additional gain in classification performance on the CALMS21 test dataset (per-frame F1 score of 0.840 ± 0.001 on foreground categories, 0.867 ± 0.001 on all categories; Figure 1C and D).

We additionally computed these F1 scores for raw model outputs, smoothed results and thresholded results on two other levels, i.e., the annotated behavioral intervals and the predicted intervals (count of behavioral interactions, SI Figure 1). We found that smoothing model outputs lowered the performance metrics on the test dataset when assessing classification performance based on annotated intervals (SI Figure 1B and C). This was likely balanced during hyperparameter optimization by a strongly increased classification performance after smoothing when focusing on the predicted intervals (as shown in SI Figure 1E and F for the test dataset). Category-specific thresholding could partially recover the annotation-based F1 scores, but lowered classification performance when focusing on predicted behavioral intervals. See SI Table 1 for detailed results.

### Social cichlids dataset

We also tested our package on a novel dataset of social interactions in groups of cichlid fish. This dataset presented a more complex scenario, requiring not only accurate classification of dyadic behaviors but also proper disentanglement of interactions among multiple individuals, each engaging with only one partner at a time. The social cichlids dataset presents further challenges due to a higher number of behavioral foreground categories (six categories, see Table 1) and through the types of annotated social behaviors. These include behavioral categories that, although distinctly defined, may overlap in spatiotemporal features. For example, the categories ‘approach’, ‘bite/dart’ and ‘chase’ all entail that the focal individual moves directedly towards a recipient fish. Threat or submissive displays (‘frontal display’, ‘lateral display’ and ‘quiver’) do not require direct contact between individuals, and are generally more inconspicuous than other categories such as ‘chase’. Nevertheless, distinct behaviors that overlap kinematically and can be performed between actors and recipients over long distances represent natural behavioral variation that is valuable to quantify.

**Table 1:** Ethogram of N. multifasciatus that was used for the behavioral scoring of the social cichlids dataset.

| Category | Description | Interpretation |
| --- | --- | --- |
| Approach | Actor moves directedly towards recipient | Neutral/affiliative |
| Lateral display | Actor positioned laterally in recipient's field of view, in proximity and with stiff body posture ('S-bend'), optionally undulating caudal fin towards recipient or with lateral movement | Display aggression |
| Frontal display | Actor in proximity to recipient, with stiff body posture ('S-bend') or puffed operculi, directed movement towards recipient | Display aggression |
| Bite/dart | Actor accelerates towards recipient ('dart'), attempts physical contact between mouth and recipient ('bite') | Aggressive |
| Chase | Actor follows recipient, both at elevated speed | Aggressive |
| Quiver | Actor quivers entire body in proximity to recipient | Submissive |

To train a model that can distinguish these behaviors from each other and from the behavioral background category ‘none’, we fitted an XGBoost classifier on the training subset of the social cichlids dataset (again, with number of boosting rounds set to 1000, following the procedure outlined in the methods section). We also optimized hyperparameters for the post-processing of classification results (temporal smoothing and thresholding). Similar as described above, we assessed classification performance with F1 scores on three levels: video frames, annotated intervals and predicted intervals. However, in contrast to the CALMS21 classification, we focused mainly on behavior counts (i.e., predicted intervals) instead of per-frame results, as annotations were much sparser in this dataset (as visualized in Figure 2A). We found that classification performance on this dataset was lower overall (see Figure 3B and C; macro F1 across all categories: 0.217 ± 0.004), but improved with both model output smoothing (0.375 ± 0.007) and thresholding (0.453 ± 0.008). Classification errors mainly occurred through false positive predictions when the true category was the behavioral background category ‘none’, and much less between the behavioral foreground categories (Figure 3A). See SI Figure 2, SI Figure 3 and SI Table 2 for additional results based on video frames and annotated intervals.

**Figure 2:**
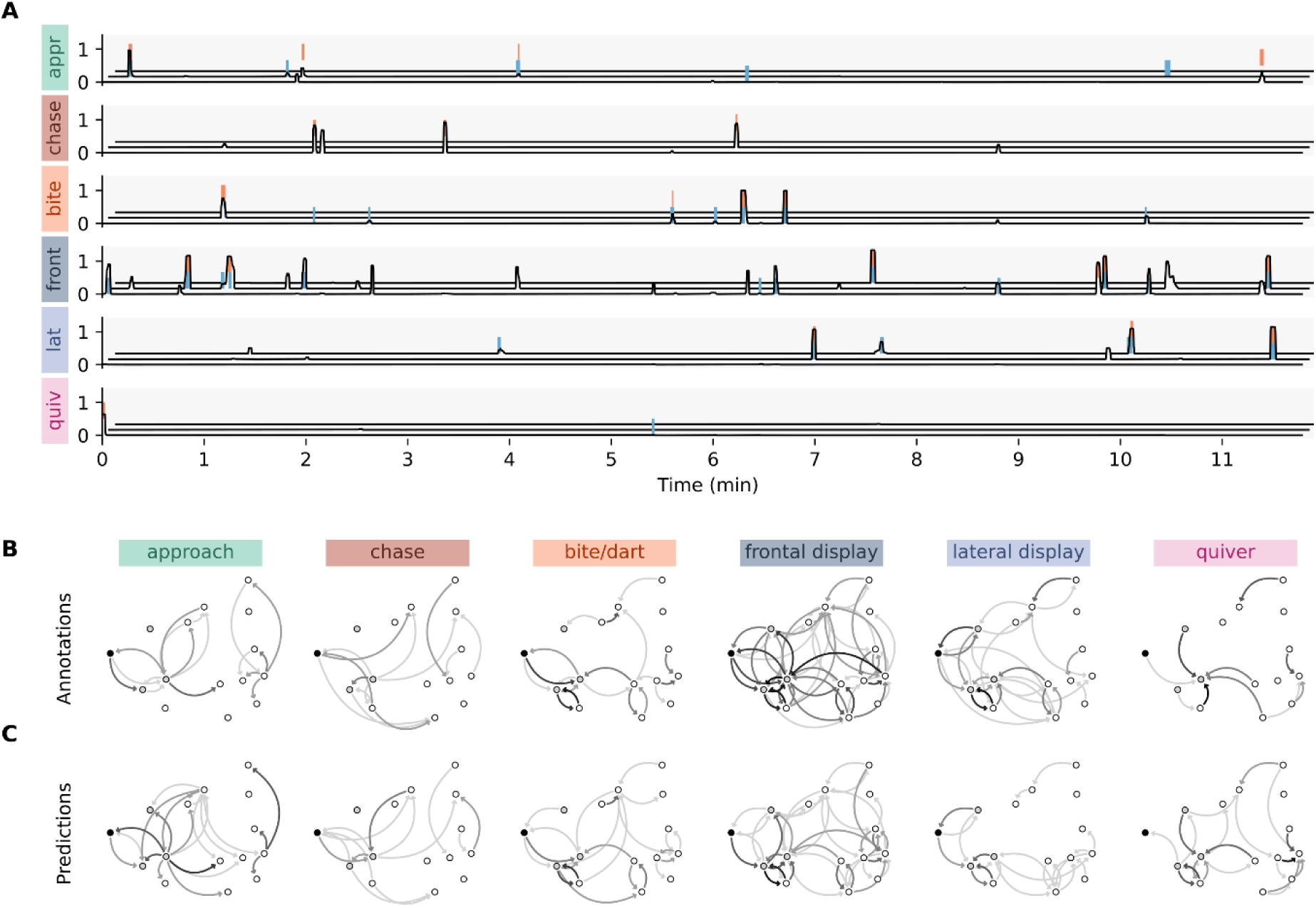
Qualitative classification results of the social cichlids dataset. **A:** Behavioral timelines for one focal fish (actor) of the test dataset and its three most frequent interaction partners (recipients). Colored bars denote ground-truth intervals (annotations, lower bars, blue) and predictions (upper bars, orange) for the six behavioral categories. Lines represent per-category model outputs (classification probabilities after smoothing). For recipient 2 and 3, intervals and lines are offset along both x and y axes. The behavioral background category was excluded for a clearer visualization with sparse behavioral data. **B and C:** Annotated and predicted interaction network of one group (15 fish), split by behavioral category. Edge line strength represents interaction counts. Note that this visualization contains data that is not part of the test dataset since the full dataset was split by individual fish and not groups. For visualization purposes, we instead used 5-fold cross validation to fit five independent classifiers that were used for predictions on the full dataset. See SI Figure 4 for correlation tests between annotated and predicted behavior counts of all dyads from the test dataset. Note: The actor and three recipients of the behavioral timelines are part of the interaction networks, marked by node color (black and grey for actor and recipients, respectively).

**Figure 3:**
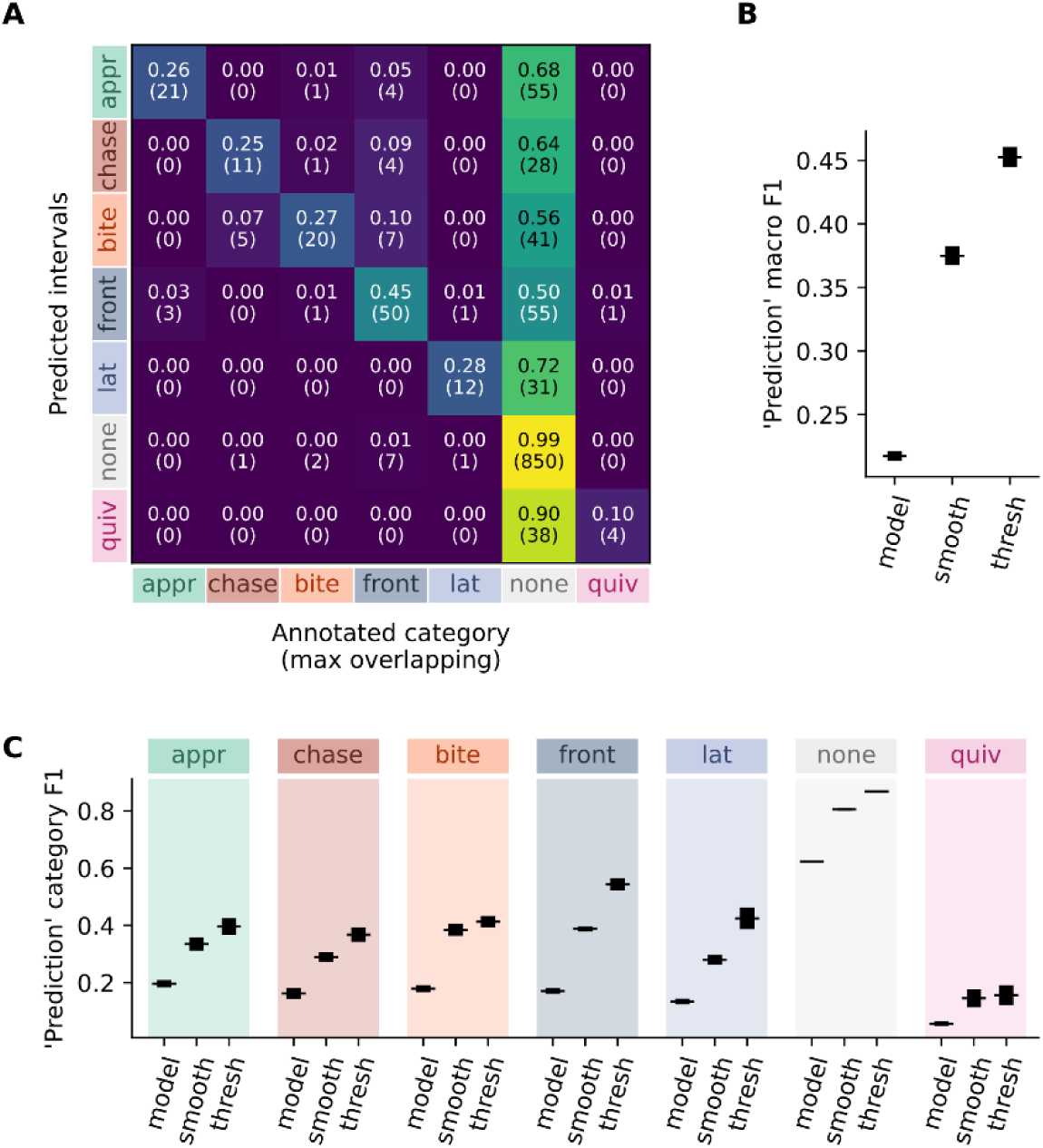
Classification results of the social cichlids test dataset. All values represent means of 20 pipeline runs with different random states, standard deviations are shown as error bars if applicable. **A:** Confusion matrix of predicted intervals and their true category (i.e., category of annotated interval with longest overlap). **B:** Unweighted average (macro) F1 score of raw model outputs (‘model’), after smoothing (‘smooth’), and after thresholding (‘thresh’), calculated across all behavioral categories on predicted intervals. **C:** Per-category F1 scores for each of the same (post-)processing steps. Note that all results were computed based on predicted behavioral intervals (i.e., interaction counts). For both video frame and annotation interval based results, see SI Figure 2 and SI Figure 3, respectively.

Despite the relatively high rates of false positive predictions in this test scenario, we found that our automated behavioral scoring pipeline could still provide otherwise inaccessible data, both in quantity (e.g., when applying classifiers on new videos) and quality. For example, interaction networks of the recorded groups based on either annotated or predicted behavior counts were visually similar (Figure 2B). To test if our model predictions were reliable proxies for observed behaviors (i.e., ground-truth annotations), we tested the correlation between these two networks for each behavioral category by correlating annotated and predicted behavior counts across all dyads of the designated test dataset. We found moderate to strong correlations between annotated and predicted counts for most categories (Pearson correlation coefficients; lowest for ‘quiver’, r = 0.19, and chase, r = 0.46, all other between 0.56 and 0.76, see SI Table 3).

Other approaches to quantify social interactions in animal groups often use behavioral proxies such as the cumulative time spent within a certain distance (e.g., association networks; Davis et al., 2018). However, these proxies cannot capture whether social interactions occur, nor the types of interactions that occur, when animals are in proximity to each other. We compared our classification method to such a non-qualitative proxy – association time, defined as the cumulative duration that individuals spent within three body lengths of one another. For this, we tested if ground-truth behavior counts correlated more strongly with counts resulting from classifier predictions than with association time, accounting for non-independence of both correlations using the William’s test with a shared variable (psych package; William Revelle, 2024; R v4.4.3; R Core Team, 2025). We found that this was the case for all six behavioral foreground categories (P < 0.001 for all categories). See SI Figure 4 for a visualizations of these correlations and SI Table 3 for statistical results and a sensitivity analysis for alternative proximity thresholds.

## Discussion

Novel methods in computational ethology have proven invaluable in describing the structure of animal behavior. With the advent of animal tracking approaches that can gather high-resolution posture and movement data in the wild (Dell et al., 2014), these methods can now be applied in contexts closer to the original goal of ethology, that is to measure the behavior of freely interacting animals in natural contexts. Here we implemented and tested the *vassi* Python package to automatically score the directed social behavior of multiple interacting animals. With our package, we were able to achieve classification results on a benchmark dataset comparable to those from existing tools (e.g., outperforming the baseline model that was published alongside the CALMS21 dataset; Sun et al., 2021). Additionally, we introduced a more naturalistic and complex test case with groups of cichlid fish (15 individuals per group). This dataset possesses properties that are commonly associated with *in-situ* observations of animals – directed interactions (i.e., incoming and outgoing social behaviors) between multiple individuals, intermittent and noisy trajectory data (e.g., fish can freely enter shelters or visually occlude each other), and continuous behavior types ranging from conspicuous aggression involving contact between individuals to subtle threat or submission displays. Testing our package on this novel dataset, we showed that the implemented behavioral classification workflow generalizes across use-cases, including different social settings (pairs versus groups) and species (mice and fish). Albeit having a lower classification performance on the cichlids data, we were able to validate the applicability of our workflow to this more complicated dataset and use the inbuilt reviewing process to verify and update ambiguous cases at need. For this, we implemented an interactive tool to inspect behavioral datasets that can be used to display video sequences of behavioral interactions, both ground-truth annotations and model predictions. Misclassifications occurred mainly between the behavioral foreground categories and the background category through false positive predictions, which may introduce systematic errors into behavioral datasets resulting from such classification workflows. However, when comparing our classification results to an alternative behavioral proxy that is commonly used for within-group behavioral interactions – association time (see e.g., Whitehead, 2008), we found that the automatically scored behaviors provide a more accurate, and importantly, a categorical representation of behavioral interactions. Proximity or association time alone, for example, are often not sufficient to capture relevant interaction networks (discussed in Farine, 2015). We further demonstrated that the post-processing routine of model output smoothing and subsequent thresholding is robust and improved classification performance on both datasets.

Our package does not pose any restrictions on the directionality of interactions or number of interacting individuals, and both tracking and behavioral data can be used without specific pre-processing strategies. Users can import individual trajectories and assign them to groups (and encompassing, datasets), which in turn can be annotated with behavioral observations from tools such as BORIS (Friard & Gamba, 2016). Since our package is primarily scripting-oriented, users can load their raw data (e.g., trajectories from a CSV file), and our package then provides all functionality that is required to train classification models on this data. Similarly, users can further apply these models to new data and score their entire datasets in an automated way. Classification results are provided in tabular form, similar to the behavioral annotations that users are familiar with. This design integrates with existing Python libraries for machine learning such as scikit-learn and XGBoost (Chen & Guestrin, 2016; Pedregosa et al., 2011), which makes the pipeline that we implemented in the *vassi* package extensible and open for future innovation in this field. For example, the process of model fitting is not hidden behind multiple layers of code but directly exposed, users can therefore easily exchange, test and evaluate different model types. Similarly, our package provides a broadly applicable feature extraction workflow with spatiotemporal features such as keypoint distances or posture angles. Although the current implementation focuses on the two case studies with data derived from 2D-tracking, this workflow can be extended to 3D data without requiring modifications of the overall classification pipeline.

*vassi* provides a compromise between accessibility (i.e., providing a fully featured graphical user interface, GUI) and extensibility or integration with other software (e.g., by providing a well-documented application programming interface, API). The package includes a widget-based GUI to facilitate interactive visualization of classification results to help improve overall classification performance through iterative validation and re-training (e.g., implemented in JAABA and A-SOiD; Kabra et al., 2013; Tillmann et al., 2024). This GUI focuses on two main aspects that become essential in more complex behavioral scenarios: First, the inspection of behavioral predictions (or annotations) across entire projects through an interactive, feature-rich table that supports filtering, editing and sorting; and second, the generation of high-quality video snippets of behavioral sequences. This interactive tool can also be used as a standalone application to make large behavioral video datasets accessible, and may help researchers to comply with best practices demanded for this research area (e.g., data transparency and sharing; Luxem et al., 2023). The remaining functionality of our package (i.e., data handling, feature extraction, model training and predictions, post-processing of classification results, hyperparameter optimization) is accessible via its API that allows users to implement their own behavioral scoring pipelines without the necessity of writing custom code.

Our package is not the first example of a behavioral classification workflow, existing tools such as JAABA, SimBA, MARS or A-SOiD implement similar methods and functionality (Kabra et al., 2013; Goodwin et al., 2024; Segalin et al., 2021; Tillmann et al., 2024). The general approach is conserved across these existing tools, using animal trajectories (here, always derived from video data) and behavioral annotations to extract spatiotemporal features serving as input to train supervised machine-learning models that can then automatically classify behaviors in further data. Although implementation details differ between these tools, they share two main limitations, firstly the number of individuals that can interact with each other in the video observations (for example, limiting to only one or two behaving individuals), and secondly, a limited perspective of directedness of behavior in dyadic animal interactions (i.e., certain behaviors can only occur in one direction between two individuals). These two limitations preclude studying social behavior as recursive responses to social stimuli between individuals, as naturally occurs in animal groups. Both limitations arise from software design, most of the tools mentioned above treat multiple animals as one ‘entity’ by combining their keypoints in a fixed order before feature computation. This order then imposes the direction in which behaviors can be classified, both MARS and A-SoiD, for example, only consider the behaviors of one individual (the resident mice in the CALMS21 dataset). With these tools, workarounds for classifying behaviors in multi-animal settings, such as reloading projects with switched trajectories, may be feasible for two individuals but become impractical as the number of tracked animals increases. Only JAABA, the oldest of the existing tools, demonstrated its utility on a dataset of animal groups (multiple fruit flies, *Drosophila melanogaster*) by presenting an alternative in which the number of individuals is not restricted, and behavioral classifiers are trained on the dyadic spatiotemporal features of a focal individual and its nearest neighbor in each video frame. This however assumes that interactions always happen between nearest neighbors, which does not generalize to all behavior types, for example threat displays that might be displayed over larger distances.

An alternative pathway for automated behavioral classification uses video data directly. The tools DeepEthogram (Bohnslav et al., 2021) and DeepAction (C. Harris et al., 2023) employ convolutional neural networks to extract relevant pixel-based features from video frames, which is then used for behavioral classification. This is an efficient end-to-end approach from videos to behavioral scoring results, however, it causes stronger limitations on the type of videos that can be analyzed when compared to the more common workflow with posture tracking and the calculation of spatiotemporal features. For example, disentangling directed behavioral interactions of two visually similar animals is not possible with this approach, but can be done when both animals are individually tracked. This is especially problematic when analyzing videos of multiple interacting animals for which the classification of directed behavioral interactions would not only require visually distinct individuals, but also that the classification model can learn a representation of each animal and uses this information for its behavioral predictions.

The *vassi* package overcomes these challenges and provides an automated behavioral scoring pipeline from animal tracking data that can be used to classify social and non-social behaviors in various contexts – single individuals, dyadic interactions, and animal groups. As such, the package broadens the applicability of methods that continue to gain importance in the subfields of neurobiology and computational ethology to other research areas of behavioral biology, particularly in cases where behaviors vary along a continuum or in which multiple animals are interacting at once. *vassi* further enables researchers to visualize and validate behavioral events, contributing to the adoption of reproducible, computational methods in behavioral studies, which may help to alleviate the replication crisis in field-based ecological research (Filazzola & Cahill Jr, 2021). Overall, this approach both identifies emerging challenges in the application of computational methods to more naturalistic settings, and provides a framework for meeting these challenges.

## Methods

### Implementation

The *vassi* package implements a complete data pipeline to automate the scoring of social behavior in animal groups. Users need to prepare raw data in the form of individual trajectories and matching behavioral annotations. This entails tracking one or more keypoints (e.g., body parts) of at least one focal individual, along with corresponding timestamps for when each individual starts and stops a behavioral event or social behavior towards another individual. While pose estimation methods like DeepLabCut or sleap (Mathis et al., 2018; Pereira et al., 2022) can provide detailed tracking information, event-logging software such as BORIS (Friard & Gamba, 2016) can be used to obtain the required behavioral data.

As outlined step-by-step below, the package provides multiple core features, including trajectory preprocessing, feature engineering and extraction, dataset sampling for classifier training or validation (e.g., splitting and k-fold for cross-validation), and post-processing of classification results. Additionally, an interactive visual inspection tool enables users to render video snippets of behavioral sequences for both raw input data (trajectories and behavioral annotations) and model predictions (see Figure 4 and SI Video 1). Refer to the online documentation at *vassi.readthedocs.io* and our GitHub repository (*github.com/pnuehrenberg/vassi*) for example scripts and two case studies.

**Figure 4:**
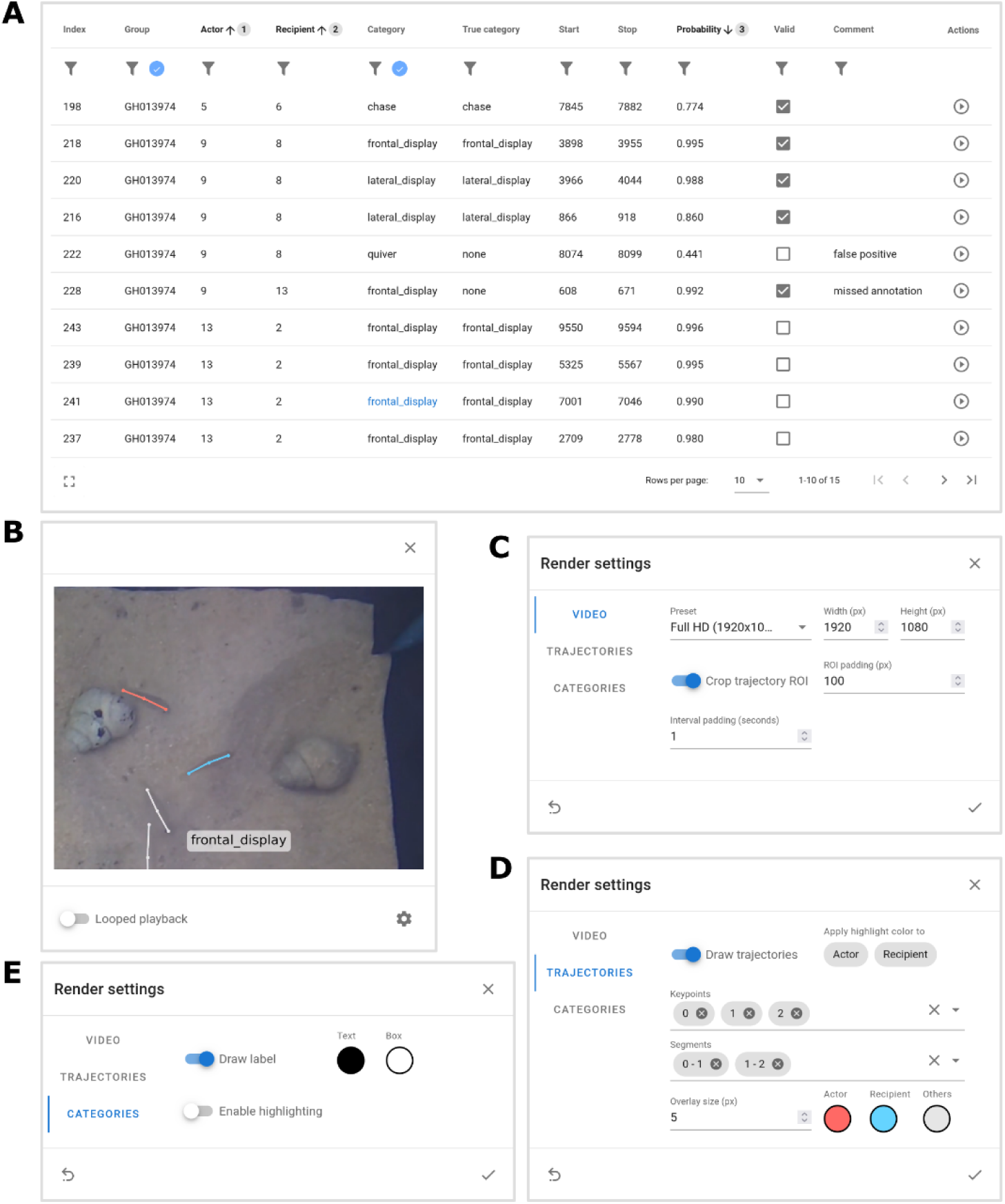
Interactive validation tool. **A:** An interactive table widget enables the inspection and editing of entire behavioral datasets from within the JupyterLab coding environment. Columns can be sorted (with multiple sorting, in the example: 1 – ‘Actor’ – ascending, 2 – ‘Recipient’ – ascending, 3 – ‘Probability’ – descending) and interactively filtered through selection, value ranges or quantile ranges (accessible via respective pop-ups). Active filters are depicted with the blue tick button. All fields are editable, but only allow valid entries if applicable (e.g., existing behavioral categories). Boolean columns are exposed as check boxes. Free text input is also possible, e.g., for comments. Action buttons can link to other widgets, as for example, video playback. **B:** Video playback interface to visualize behavioral sequences. Each row in the interactive table (i.e., annotations or predictions depending on use case) can be used to render a corresponding video, optionally with overlays for tracking data and a behavioral label. Videos are played directly in the interface and can be looped, stopped and maximized. **C – E:** Rendering options to configure video output. In the example, both actor and recipient are highlighted with red and blue, respectively.

### Handling trajectory data

Our package provides functionality to handle and process trajectory data. For example, users can load and save of trajectory data from or to structured HDF5 files (a standard for multidimensional numerical data; The HDF Group, n.d.). Data access and manipulation is facilitated through object-oriented Python structures that provide methods such as temporal resampling, interpolation (e.g., to impute missing tracking data for occluded individuals) or time-window slicing (e.g., to select relevant intervals or temporally align multiple trajectories). These trajectory objects also allow direct data access, implementing common indexing operations similar to the widely-used Python packages NumPy and pandas (C. R. Harris et al., 2020; The pandas development team, 2024).

### Feature engineering and extraction

Our package offers a user-friendly workflow to calculate spatiotemporal features from individual trajectories or dyads (i.e., ordered trajectory pairs). Features such as ‘posture angles’, ‘keypoint distances’, ‘speed’, or ‘target velocity’ can be configured through a single YAML file (a human-friendly data serialization language), allowing users to reproduce their feature extraction workflows across different datasets or studies. The features are implemented as Python functions with a common set of inputs. For example, ‘keypoint distances’ can be used to calculate within-animal keypoint distances if only one trajectory is specified, or alternatively to calculate between-animal distances with two trajectories specified. Users can define these features as either ‘individual’ or ‘dyadic’ in their configuration and only need to provide the relevant keypoints or keypoint pairs. In the case of temporal features, such as ‘speed’ (i.e., displacement over time), users can specify a temporal step over which the feature is calculated. Feature extraction workflows can be further augmented with data transformations such as standardization or discretization, and are compatible with data transformations implemented in the scikit-learn library (Pedregosa et al., 2011). Our package additionally implements a time-window transformation to compute sliding time-window representations of the extracted features, allowing workflows similar to JAABA or MARS (Kabra et al., 2013; Segalin et al., 2021)

### Training classifiers, prediction and post-processing

To simplify handling entire datasets that combine tracking data with behavioral annotations, our package provides functionality through Python objects for common operations such as data validation, sampling, or subsampling. These objects reflect the inherent structure of such datasets: ‘Groups’ are initialized with trajectories and focus on either ‘individuals’ (for non-social behavior) or ‘dyads’ (for social behavior), a ‘dataset’ consists of at least one of such groups. Users can annotate each of these data objects with intervals of one or more behavioral categories. Using the feature extraction workflow described above, sampling annotated datasets results in a feature table with corresponding target categories, while sampling unannotated datasets returns feature tables alone, suitable for the classification of new data or for unsupervised learning. This design integrates seamlessly with machine learning libraries like scikit-learn or XGBoost (Pedregosa et al., 2011; Chen & Guestrin, 2016), allowing users to train classification models without writing custom code.

Our package enables users to directly apply such classification models on entire datasets or their subcomponents (i.e., groups, individuals, or dyads). This prediction step is made accessible to users through a common code interface: raw model outputs (per-frame category probabilities) are transformed into behavioral predictions by grouping consecutive frames of the same category into intervals. The resulting data is provided in tabular form as pandas DataFrames and can be directly used in down-stream statistical analyses.

Additionally, users can apply two post-processing steps to classification results. Model outputs can be temporally smoothed by applying one or more smoothing methods (for example filters implemented in SciPy’s signal module; Virtanen et al., 2020), reducing noise in the time-series of category probabilities. This may help to reduce the number of short behavioral predictions and limits the amount of frequent back-and-forth transitions between predicted behavioral categories. In addition, users can specify a probability threshold for each category, only keeping predictions above their category’s respective threshold.

### Optimization of hyperparameters

Predictive performance in machine learning often depends on hyperparameters that define model structure and optimization routines, for example decision tree depth or learning rate (Yang & Shami, 2020). Pre- and postprocessing workflows can also introduce hyperparameters, such as category-specific subsampling frequencies or decision thresholds for classification outputs (Sauer et al., 2024).

With the *vassi* package, we provide methods to optimize the parameters arising from the two implemented post-processing steps: (I) the amount or duration of temporal smoothing to reduce temporal noise in classification results, and (II) category-specific decision thresholds. For binary classification, this threshold typically defaults to 0.5, while in multi-class tasks the category with the highest probability is selected. However, such general rules may not be optimal for all datasets. Our package enables users to tune these hyperparameters with optuna (a hyperparameter optimization framework; Akiba et al., 2019) to maximize predictive performance with regard to their data and use case.

### Model validation

Users can choose any classification model compatible with the scikit-learn classification API (Pedregosa et al., 2011). Therefore, comparing classification results across models can help to identify the best-performing model for a given dataset or guide the selection of post-processing routines to improve performance. In classification tasks, recall and precision are key metrics for model evaluation (Juba & Le, 2019). Recall measures the proportion of true positives correctly identified for a category at a given decision threshold, while precision reflects the proportion of predicted positives that genuinely belong to the target category. Both metrics depend on the decision threshold and are interacting through the relationship between true positive and false positive predictions.

A commonly used composite metric is the F1 score, the harmonic mean of precision and recall. Certain behavioral categories that occur only sparsely in a behavioral dataset may be as critical to classify accurately as other, more common categories. Therefore, the macro F1 score (an unweighted average across categories) can be particularly useful to evaluate classification with imbalanced datasets (e.g., Sun et al., 2021; Tillmann et al., 2024). Our package provides methods to compute this macro F1 score for test data on three different levels: (I) per-frame, assessing classification performance based on the cumulative duration of each behavioral category, (II) on annotated behavioral intervals, focusing on the recall of ground-truth intervals, and (III), on predicted behavioral intervals, evaluating the precision of predicted intervals. The two interval-based scores measure performance while considering interval counts rather than cumulative durations.

Another tool for assessing classification performance are confusion matrices, showing true and false predictions and the amount of misclassification between categories. Our package implements methods to compute confusion matrices directly on the classification results of annotated data. Equivalently to the F1 score, confusion matrices can be generated on three levels, per-frame, on annotation intervals, and on prediction intervals.

### Visual inspection of behavioral sequences

We provide users with two visual inspection tools for behavioral sequences. The first is a timeline-plot (Gantt chart) to render behavioral sequences for one individual or dyad at a time, also implemented for example in BORIS, SimBA or TIBA (Friard & Gamba, 2016; Goodwin et al., 2024; Kraus et al., 2024). Users can augment this visualization with raw and smoothed model outputs (i.e., classification probabilities for each class) and choose whether to include predicted or annotated intervals, or both.

The second inspection tool is a feature-rich table that enables data filtering, sorting and editing and can be used to render video snippets of behavioral sequences (Figure 4). Each row in this table describes one behavioral interval, either annotation or prediction, depending on the input. The interactivity allows users to inspect entire datasets, for example, by filtering intervals by category (e.g., showing 2 out of 6 categories), and out of these only keep the bottom 10% by classification performance (applying a conditional quantile filter). A workflow such as this enables efficient validation of behavioral predictions, while, in the example case, focusing on likely misclassifications. Users can add columns to the table, for instance for comments or Boolean indicators (e.g., to mark if a row is considered valid or rejected). From each row, a widget can be opened to render the correspondent video sequence. We enable users to adjust options of the graphical display of these video sequences, for example by selecting the tracked keypoints or segments to visualize, the rendering resolution or cropping the video around actor and recipient individuals. See *SI Video 1* for an exemplary use of this tool.

### Case studies

We tested the *vassi* package in two case studies, firstly with the CALMS21 benchmark dataset of mouse resident-intruder interactions (Sun et al., 2021), and secondly with a more complex dataset of social behavior in groups of cichlid fish. In both cases, we fitted an XGBoost classifier to distinguish between behavioral categories in dyadic social interactions. These categories included defined social behaviors directed from an actor to a recipient (foreground categories) and a background category encompassing all other behaviors.

### CALMS21 dataset

The CALMS21 dataset supplies benchmarks for multiple tasks, but we focused solely on the first task ‘classical classification’. This benchmark comprises 89 resident-intruder video sequences, with 19 designated as the test set. We adhered to this split, using the first 70 sequences for model training and hyperparameter tuning. Each video frame is categorized into one of four behavioral categories: three social behaviors (‘attack’, ‘mount,’ and ‘investigation’) as foreground categories, and a fourth (‘other’) as the background category assigned to unlabeled frames. Interactions are only one-directional, from resident to intruder. The dataset also includes posture tracking data for resident and intruder mice, with seven body parts tracked per mouse (keypoints on nose, left and right ear, back of neck, left and right hip, and base of tail). Both the behavioral annotations and tracking data were used without modification.

The CALMS21 dataset contains imbalanced categories (see SI Table 4) and large datasets can pose computational challenges. To address both, we randomly subsampled the overrepresented categories (‘investigation’ and ‘other’) during classifier training, limited to 30000 samples each. Subsampling was stratified by behavioral intervals to maintain proportional representation of all behavioral annotations. SI Table 4 provides an overview of category frequencies in the training data.

During data sampling, we computed movement and posture features for each selected video frame (see SI Table 6 for a full list). Using this data, we trained an XGBoost classifier with the number of boosting rounds set to 1000 to allow for a more complex model but otherwise used default parameters. We used category-specific sample weights via the scikit-learn method ‘compute_sample_weight’ to further mitigate imbalanced categories. The fitted classifier was then applied to the 19 video sequences of the CALMS21 test dataset to evaluate the performance of our package’s classification pipeline. The model generated raw output in the form of classification probabilities for each category across all frames of all sequences. In post-processing, we selected the category with the highest probability for each frame and grouped consecutive frames of the same category into behavioral intervals. These intervals were arranged in a table with columns for ‘group’ (i.e., sequence ID), ‘category’, ‘start’, and ‘stop’, the endpoint of behavioral results in the *vassi* package.

We calculated the per-frame macro F1 score for the three foreground categories to enable comparisons with Sun et al. (2021). However, we also deemed the background category’s accuracy important for model evaluation and therefore additionally calculated the per-frame macro F1 score with all four categories. We further computed two interval-based F1 scores: one focusing on the recall of annotations (i.e., how many annotation intervals were overlapping primarily with predictions of the same category) and another assessing the precision of predictions (i.e., how many predicted intervals had the same category as the annotation with highest temporal overlap). We also evaluated model performance with confusion matrices, both on the per-frame level and on annotated and predicted intervals.

Subsequently, we tuned two sets of post-processing hyperparameters to improve classification results. We sought to find the optimal duration of output smoothing when applying a filter (e.g., mean or median) to classification probabilities, and secondly, the optimal decision threshold for each category. We used k-fold cross-validation (k = 5) on the original CALMS21 training dataset, splitting the dataset so that each of the 70 video sequences was used for validation in exactly one of the five dataset folds. For each fold, we sampled training data using the same features and category frequencies as before (see SI Table 4) to fit a total of five XGBoost classifiers. These classifiers were then used for prediction on the respective test data of each fold. Subsequently, we used optuna (Akiba et al., 2019), a hyperparameter optimization framework, to tune the specified parameters within a predefined search space. Finally, we applied these two optimized post-processing steps to all predictions on the CALMS21 test dataset, and compared the results to the baseline without post-processing and the results from Sun et al. (2021).

### Social cichlids dataset

We further tested the *vassi* package with a more complex dataset of socially interacting cichlid fish. This dataset was collected as part of a larger study (Bose et al., in prep) on *Neolamprologus multifasciatus*, a shell-dwelling cichlid fish of Lake Tanganyika that lives in social groups and exhibits a range of social behaviors (Kohler, 1998).

We selected nine videos of captive *N. multifasciatus* groups of 15 individuals each, filmed top-down with a GoPro Hero7 action camera at a resolution of 2704 × 1520 pixels and 30 frames per second. The video recordings captured the aquarium setups that housed the groups of fish during the experiments. These aquaria were enriched with sand as bottom substrate and 30 empty *Neothauma tanganyicense* snail shells as shelters. As part of the overarching study, we tracked individual fish in these videos using a custom object detection and posture estimation workflow (using the detectron2 framework; Wu et al., 2019; and Python scripts and user interfaces adapted from Francisco et al., 2020). As a result, each fish was tracked with three keypoints along the main body axis (head, dorsal center, tail) when visible and detected by the detectron2 model. For all other frames, the keypoints were linearly interpolated between the previous and the next detection. Additionally, one experimenter scored the social behaviors of all fish (‘approach’, ‘lateral display’, ‘frontal display’, ‘bite/dart, ‘chase’, ‘quiver’ as foreground categories, see Table 1 for descriptions) using the event logging software BORIS (Friard & Gamba, 2016), always in relation to a recipient individual. The trajectories and behavioral annotations are made available alongside the *vassi* package and were used without further modification.

We followed the same workflow as with the CALMS21 dataset but implemented changes where differences in dataset structure demanded so. For example, we allowed interactions between individuals to be bi-directional, which resulted in 210 directed dyads for group each group of 15 individuals (1862 dyads across the nine videos, one group had only 14 individuals). Out of these dyads, we randomly selected 20% of individuals as actors (i.e., the first individual in a directed dyad; 375 dyads) for testing, the remaining 1487 dyads were used for data sampling and the training of classifiers. Similar to the CALMS21 dataset, our social cichlids dataset also consisted of imbalanced behavioral categories, and we chose to subsample the overrepresented categories ‘none’ and ‘frontal display’ for training classifiers (see SI Table 5). Note that since each individual could interact with all others in its group, but only with one at a time, the background category ‘none’ was substantially larger in this dataset. To address this, in addition to random sampling across all ‘none’ samples, we also selected samples where an individual showed a behavior towards a recipient fish, but then specifically sampled another of its close neighbors (to whom the behavior was not directed) to obtain samples that might be difficult to classify. The number of tracked keypoints (three keypoints, see above) also differed from the CALMS21 dataset, hence, the composition and number of extracted features were different as well. Since the posture information of the tracked fish was less detailed than in case of the CALMS21 dataset, we further transformed the features using a sliding time-window aggregation on a 3 second timescale (93 video frames) with three consecutive windows (first, second and third second), for which descriptive statistics such as the mean, median and quantile values were computed.

After fitting a XGBoost classifier on the training data of the social cichlids dataset, we also optimized two post-processing hyperparameters (temporal smoothing and decision thresholds). Here, instead of splitting the training dataset by videos during 5-fold cross-validation, we split groups by dyads, the smallest behaviorally independent element in this dataset. Note that this was in fact equivalently implemented to the cross-validation splits for the CALMS21 dataset, as video sequences in the CALMS21 dataset consist of ‘groups’ containing exactly one directed dyad (i.e., resident to intruder). After the optimal values of these hyperparameters were determined for the social cichlids dataset, we applied both post-processing steps with these values on the social cichlids test dataset. Classification performance was evaluated with F1 scores (as described before) and compared between the baseline without post-processing and results with temporal smoothing and adjusted decision rules.

### Software development

We developed the *vassi* package in Python 3 (version >=3.12), building on established methods for data handling and manipulation (NumPy, pandas, h5py and SciPy), and machine learning packages (scikit-learn, XGBoost and optuna). The package was implemented with a focus on interactive coding in the JupyterLab environment. All steps of the behavioral scoring pipeline are exposed in an object-oriented way through classes and their respective methods. The inspection tool for model validation was implemented with the ipywidgets and ipyvuetify packages. All components of the *vassi* package can be installed with a single command and across operating systems (tested on Windows 11, Ubuntu 24.04.2 and macOS Sequoia 15.3.1). The documentation of the package and example scripts are available online at *vassi.readthedocs.io*.

Software development and tests were conducted on consumer-grade hardware, a laptop with an Intel Core i7-1165G7 CPU and 16 GB of RAM (for both Windows and Ubuntu), a workstation with an Intel Core i9-9900X CPU and 64 GB of RAM (Ubuntu) and a MacBook Pro 2024 with an Apple M4 Pro chip and 24 GB of RAM (macOS). Hyperparameter tuning was performed on the computing cluster ‘Raven’ of the Max Planck Computing and Data Facility.

## Code and data availability

All code and data is open-source and publicly available. The *vassi* package is under version control on GitHub at *github.com/pnuehrenberg/vassi*, example scripts (including scripts to reproduce all results and figures) and code documentation are also available online (*vassi.readthedocs.io*). The social cichlids dataset (videos, trajectories and behavioral annotations), all source files of the *vassi* package, and all additional data or configuration files can be accessed at https://doi.org/10.17617/3.3R0QYI.

## Ethics

We did not conduct any animal experiments that were designed for the purpose of creating the presented methodology. The two case studies were performed on either existing data (CALMS21 mice dataset; Sun et al., 2021) or on data collected for an overarching study on cichlid fish that was conducted in accordance with the Directive 2010/63/EU on the protection of animals used for scientific purposes and the Tierschutzgesetz (TierSchG) Baden-Württemberg, approved by permit no. G 19/79.

## Acknowledgements

We thank all members of the Behavioral Evolution Research Group for feedback and valuable discussion. This research was funded by the Deutsche Forschungsgemeinschaft (DFG, German Research Foundation) under Germany’s Excellence Strategy – EXC 2117 – 422037984, by the Max Planck Institute of Animal Behavior and Vetenskapsrådet (grant number 2023-03866 to AB).

## Author contributions

**PN:** Conceptualization, Methodology, Software, Validation, Formal Analysis, Data Curation, Writing – Original Draft, Writing - Review & Editing, Visualization. **AB:** Conceptualization, Validation, Resources, Data Curation, Writing - Review & Editing. **AJ:** Conceptualization, Resources, Writing - Review & Editing, Supervision, Funding acquisition.

## Supplementary Information

### Supplementary Tables

**SI Table 1:** CALMS21 test dataset evaluation results. The values report means and standard deviations of 20 pipeline runs with different random states. F1 scores were calculated on three levels, i.e., frames, annotated intervals and predicted intervals (both evaluated as counts), and for three processing steps (raw model outputs, classification results after probability smoothing and after thresholding). Classification performance was evaluated separately for each behavioral category and as unweighted average (Macro F1) across all categories and across the three behavioral foreground categories ‘attack’, ‘investigation’, and ‘mount’. All scores are reported as means and standard deviations of 20 pipeline runs with different random states.

| Level | Classification step | Macro F1 |  | Category F1 |  |  |  |
| --- | --- | --- | --- | --- | --- | --- | --- |
|  |  | foreground | all | Attack | Investigation | Mount | Other |
| Frames | Raw model outputs | 0.804±0.001 | 0.840±0.001 | 0.664±0.003 | 0.832±0.001 | 0.915±0.001 | 0.949±0.000 |
|  | Smoothed results | 0.838±0.001 | 0.866±0.001 | 0.752±0.002 | 0.843±0.001 | 0.920±0.001 | 0.949±0.000 |
|  | Thresholded results | 0.840±0.001 | 0.867±0.001 | 0.753±0.003 | 0.843±0.001 | 0.924±0.001 | 0.949±0.000 |
| Annotated intervals | Raw model outputs | 0.778±0.004 | 0.802±0.003 | 0.632±0.007 | 0.802±0.004 | 0.899±0.006 | 0.877±0.004 |
|  | Smoothed results | 0.665±0.004 | 0.695±0.003 | 0.562±0.005 | 0.675±0.005 | 0.758±0.005 | 0.785±0.004 |
|  | Thresholded results | 0.678±0.003 | 0.704±0.003 | 0.560±0.004 | 0.690±0.004 | 0.785±0.005 | 0.782±0.004 |
| Predicted intervals | Raw model outputs | 0.479±0.002 | 0.510±0.002 | 0.462±0.004 | 0.520±0.004 | 0.455±0.005 | 0.604±0.004 |
|  | Smoothed results | 0.864±0.005 | 0.879±0.004 | 0.812±0.010 | 0.866±0.004 | 0.914±0.009 | 0.923±0.003 |
|  | Thresholded results | 0.858±0.006 | 0.874±0.005 | 0.812±0.012 | 0.864±0.004 | 0.899±0.008 | 0.919±0.002 |

**SI Table 2:** Social cichlids test dataset evaluation results. All values report means and standard deviations of 20 pipeline runs with different random states. F1 scores were calculated on three levels, i.e., frames, annotated intervals and predicted intervals (both evaluated as counts), and for three processing steps (raw model outputs, classification results after probability smoothing and after thresholding). Classification performance was evaluated separately for each behavioral category and as unweighted average (Macro F1) across all categories and across the six behavioral foreground categories (excluding ‘none’).

| Level | Classification step | Macro F1 |  | Category F1 |  |  |  |  |  |  |
| --- | --- | --- | --- | --- | --- | --- | --- | --- | --- | --- |
|  |  | fore-ground | all | Approach | Chase | Bite/dart | Frontal display | Lateral display | None | Quiver |
| Frames | Raw model outputs | 0.247<br>±0.003 | 0.354<br>±0.003 | 0.237<br>±0.010 | 0.315<br>±0.007 | 0.298<br>±0.007 | 0.317<br>±0.004 | 0.219<br>±0.004 | 0.995<br>±0.000 | 0.095<br>±0.008 |
|  | Smoothed results | 0.271<br>±0.005 | 0.374<br>±0.005 | 0.262<br>±0.017 | 0.337<br>±0.008 | 0.327<br>±0.010 | 0.353<br>±0.008 | 0.242<br>±0.009 | 0.996<br>±0.000 | 0.101<br>±0.019 |
|  | Thresholded results | 0.289<br>±0.005 | 0.390<br>±0.004 | 0.283<br>±0.017 | 0.378<br>±0.010 | 0.316<br>±0.009 | 0.394<br>±0.012 | 0.258<br>±0.020 | 0.997<br>±0.000 | 0.102<br>±0.021 |
| Annotated intervals | Raw model outputs | 0.552<br>±0.013 | 0.605<br>±0.012 | 0.495<br>±0.035 | 0.535<br>±0.014 | 0.614<br>±0.022 | 0.674<br>±0.014 | 0.586<br>±0.031 | 0.919<br>±0.003 | 0.409<br>±0.069 |
|  | Smoothed results | 0.521<br>±0.012 | 0.576<br>±0.010 | 0.487<br>±0.032 | 0.537<br>±0.016 | 0.550<br>±0.012 | 0.664<br>±0.011 | 0.571<br>±0.024 | 0.907<br>±0.003 | 0.318<br>±0.058 |
|  | Thresholded results | 0.482<br>±0.015 | 0.536<br>±0.013 | 0.542<br>±0.035 | 0.557<br>±0.024 | 0.524<br>±0.015 | 0.524<br>±0.020 | 0.426<br>±0.031 | 0.864<br>±0.003 | 0.316<br>±0.064 |
| Predicted intervals | Raw model outputs | 0.150<br>±0.005 | 0.217<br>±0.004 | 0.196<br>±0.013 | 0.162<br>±0.020 | 0.179<br>±0.012 | 0.171<br>±0.010 | 0.134<br>±0.010 | 0.623<br>±0.002 | 0.057<br>±0.008 |
|  | Smoothed results | 0.303<br>±0.008 | 0.375<br>±0.007 | 0.335<br>±0.023 | 0.290<br>±0.017 | 0.385<br>±0.021 | 0.387<br>±0.009 | 0.279<br>±0.017 | 0.805<br>±0.004 | 0.145<br>±0.031 |
|  | Thresholded results | 0.383<br>±0.009 | 0.453<br>±0.008 | 0.396<br>±0.029 | 0.368<br>±0.025 | 0.413<br>±0.020 | 0.543<br>±0.021 | 0.424<br>±0.038 | 0.868<br>±0.003 | 0.156<br>±0.035 |

**SI Table 3:** Correlation tests between correlations of ground-truth behavioral interaction counts (annotations) and two potential proxies: (1) predicted counts as resulting from the classification pipeline and, (2) association time, the cumulative duration that two individuals spend within a defined distance threshold (3 average body length, and 1 and 5 body length as a sensitivity analysis). We used Williams’ tests between two dependent correlations that share one variable (two tailed) to assess whether correlations with predicted counts were different in strength compared to equivalent correlations with association time. Note that the correlation coefficients with predicted counts are repeated for each association threshold.

| Association threshold<br>(body lengths) | Category | Pearson correlation coefficient |  | Williams' Test |  |
| --- | --- | --- | --- | --- | --- |
|  |  | of annotated interaction counts<br>~ predicted counts | ~ association time | Test statistic (t) | P-value |
| 1 | Approach | 0.655 | 0.310 | 14.399 | < 0.001 |
|  | Chase | 0.463 | 0.100 | 10.137 | < 0.001 |
|  | Bite/dart | 0.759 | 0.411 | 22.391 | < 0.001 |
|  | Frontal display | 0.637 | 0.248 | 18.153 | < 0.001 |
|  | Lateral display | 0.564 | 0.254 | 12.390 | < 0.001 |
|  | Quiver | 0.189 | 0.050 | 4.777 | < 0.001 |
| 3 | Approach | 0.655 | 0.260 | 14.960 | < 0.001 |
|  | Chase | 0.463 | 0.044 | 11.499 | < 0.001 |
|  | Bite/dart | 0.759 | 0.263 | 24.686 | < 0.001 |
|  | Frontal display | 0.637 | 0.275 | 19.408 | < 0.001 |
|  | Lateral display | 0.564 | 0.268 | 11.566 | < 0.001 |
|  | Quiver | 0.189 | 0.068 | 3.839 | < 0.001 |
| 5 | Approach | 0.655 | 0.236 | 15.423 | < 0.001 |
|  | Chase | 0.463 | 0.065 | 10.826 | < 0.001 |
|  | Bite/dart | 0.759 | 0.232 | 23.688 | < 0.001 |
|  | Frontal display | 0.637 | 0.317 | 16.138 | < 0.001 |
|  | Lateral display | 0.564 | 0.280 | 10.316 | < 0.001 |
|  | Quiver | 0.189 | 0.082 | 3.128 | 0.002 |

**SI Table 4:** Overview of sample counts (video frames) and proportions by category in the CALMS21 training and test dataset and after subsampling for model training. The ‘sampling frequency’ column shows category specific (sub-)sampling strategies employed for this dataset: full sampling (value of 1.0, all frames) and sampling with a target count (for both ‘investigation’ and ‘other’, proportion in parentheses is the realized subsampling frequency). The resulting samples that were used for model training (‘subsampled training data’) can differ from the target count due to constrains from stratified sampling by behavioral intervals.

| Category | CALMS21 training dataset | Sampling frequency | Subsampled training data | CALMS21 test dataset |
| --- | --- | --- | --- | --- |
| Attack | 14039 (2.8%) | 1.0 | 14035 (13.7%) | 12630 (4.8%) |
| Investigation | 146615 (28.9%) | 30000 (0.204) | 29999 (29.2%) | 61275 (23.4%) |
| Mount | 28615 (5.6%) | 1.0 | 28608 (27.9%) | 31848 (12.2%) |
| Other | 318469 (62.7%) | 30000 (0.094) | 29995 (29.2%) | 156354 (59.7%) |

**SI Table 5:** Overview of category counts (video frames per dyad) and proportions by category in the full social cichlids dataset, when split into training dataset (selecting all dyads where a subset of 80% of all individuals where ‘actors’) and test dataset (the remaining dyads), and after subsampling for model training. The categories were either fully sampled (sampling frequency of 1.0), or subsampled with a given frequency. Note that for the behavioral background category ‘none’, we first randomly selected 1% of all available samples (asterisk in table), and then added more samples where an actor was interacting with a different individual. For these additional ‘none’ samples, we sampled the first and second non-interacting closest neighbors with 0.1x the sampling frequency of the actor’s actual interaction, and the third, fourth and fifth neighbors with 0.05x the sampling frequency. Also note that the category ‘none’ contains more than 99% of all samples because individuals can only interact with one other at a time, but groups consist of 15 individuals.

| Category | Full dataset | Training dataset | Sampling frequency | Subsampled training data | Test dataset |
| --- | --- | --- | --- | --- | --- |
| Approach | 7699 (0.03%) | 4769 (0.02%) | 1.0 | 4769 (1.85%) | 2930 (0.06%) |
| Chase | 7412 (0.03%) | 5546 (0.03%) | 1.0 | 5546 (2.15%) | 1866 (0.04%) |
| Bite/dart | 12318 (0.05%) | 9840 (0.05%) | 1.0 | 9810 (3.81%) | 2478 (0.05%) |
| Frontal display | 39569 (0.16%) | 31663 (0.16%) | 0.25 | 7915 (3.07%) | 7906 (0.16%) |
| Lateral display | 14179 (0.06%) | 10946 (0.05%) | 1.0 | 10946 (4.25%) | 3233 (0.06%) |
| None | 24910846 (99.64%) | 19870951 (99.64%) | 0.01* | 209078 (81.22%) | 5039895 (99.62%) |
| Quiver | 10053 (0.04%) | 9344 (0.05%) | 1.0 | 9344 (3.63%) | 709 (0.01%) |

**SI Table 6:** Spatiotemporal features that were extracted to train classifiers for the CALMS21 dataset. In total, 201 feature values were extracted to describe the spatiotemporal movement patterns of dyadic mice interactions. Features are either individual or dyadic and result in at least one value per feature and combination of postural elements (i.e., keypoints or segments). Some features are temporal and were calculated for different time steps (video frames). For example, ‘speed’ (an individual feature) was calculated for three keypoints and for three steps, resulting in a total of 9 values. In comparison, ‘target velocity’ is a dyadic feature and was calculated for two keypoints of the actor mouse and along six target vectors between keypoints of the actor and recipient mouse. In comparison to ‘speed’ (a scalar feature with one value), ‘target velocity’ itself is a vector of two components, i.e., the projection and rejection of the keypoint displacement onto a target vector. For the three temporal steps, this results in 3 × 2 × 6 × 2 = 72 values.

| Count | Feature name | Feature category | Postural elements | Postural target elements | Values and modifiers | Steps (frames) |
| --- | --- | --- | --- | --- | --- | --- |
| 9 | Speed | Individual | 3 keypoints<br>• nose<br>• neck<br>• tail |  |  | • 3<br>• 15<br>• 30 |
| 42 | Projected velocity | Individual | 7 keypoints (all) | 1 orientation vector<br>• tail → neck | 2 values<br>• projection<br>• rejection (absolute) | • 3<br>• 15<br>• 30 |
| 6 | Orientation change | Individual | 2 segments<br>• neck → nose<br>• tail → neck |  | Absolute | • 3<br>• 15<br>• 30 |
| 5 | Posture angle | Individual | 5 segments<br>• tail → neck<br>• neck → left ear<br>• neck → right ear<br>• tail → left hip<br>• tail → right hip | 5 segments (paired)<br>• neck → nose<br>• neck → nose<br>• neck → nose<br>• tail → neck<br>• tail → neck |  |  |
| 72 | Target velocity | Dyadic | 2 keypoints<br>• nose<br>• neck | 6 target vectors (between individuals, actor → recipient)<br>• nose → nose<br>• nose → neck<br>• nose → tail<br>• neck → nose<br>• neck → neck<br>• neck → tail | 2 values<br>• projection<br>• rejection (absolute) | • 3<br>• 15<br>• 30 |
| 49 | Keypoint distance | Dyadic | 7 keypoints (all) | 7 keypoints (all, actor to recipient) |  |  |
| 14 | Target angle | Dyadic | 2 segments<br>• neck → nose<br>• tail → neck | 7 keypoints (all, actor segment to recipient keypoint) | Absolute |  |
| 4 | Posture alignment | Dyadic | 2 segments<br>• neck → nose<br>• tail → neck | 2 segments<br>• neck → nose<br>• tail → neck |  |  |

### Supplementary Figures

**SI Figure 1:**
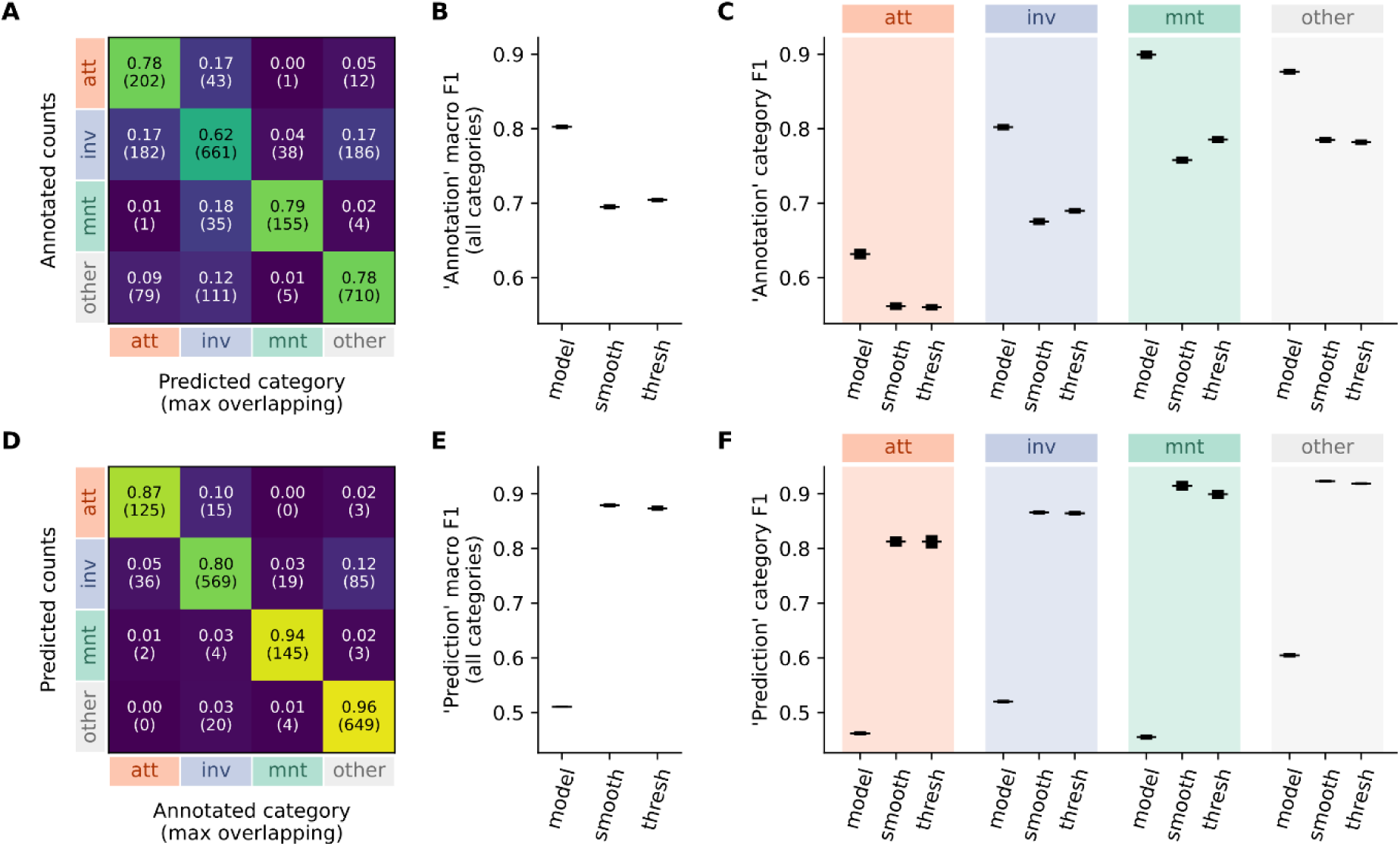
Additional CALMS21 test dataset validation results. All values represent means of 20 pipeline runs with different random states, standard deviations are shown as error bars if applicable. **A – D:** Evaluation based on annotated behavioral intervals. **A:** Confusion matrix for annotated intervals and their predicted category (i.e., category of predicted interval with longest overlap). Proportional values are normalized across rows, absolute counts are shown below in parentheses. **B:** Macro F1 scores calculated on the annotated intervals across all categories for three (post-)processing steps – raw model outputs (‘model’), after probability smoothing (‘smooth’) and after thresholding (‘thresh’). **C:** F1 scores calculated for each category and the three processing steps. **D – E:** Corresponding evaluation based on predicted behavioral intervals. **D:** Confusion matrix for predicted intervals and their true category (i.e., category of annotated interval with longest overlap). **E and F:** Macro F1 and category F1 scores calculated on predicted intervals. Behavioral categories are abbreviated: ‘attack’ – ‘att’, ‘investigation’ – ‘inv’ and ‘mount’ – ‘mnt’.

**SI Figure 2:**
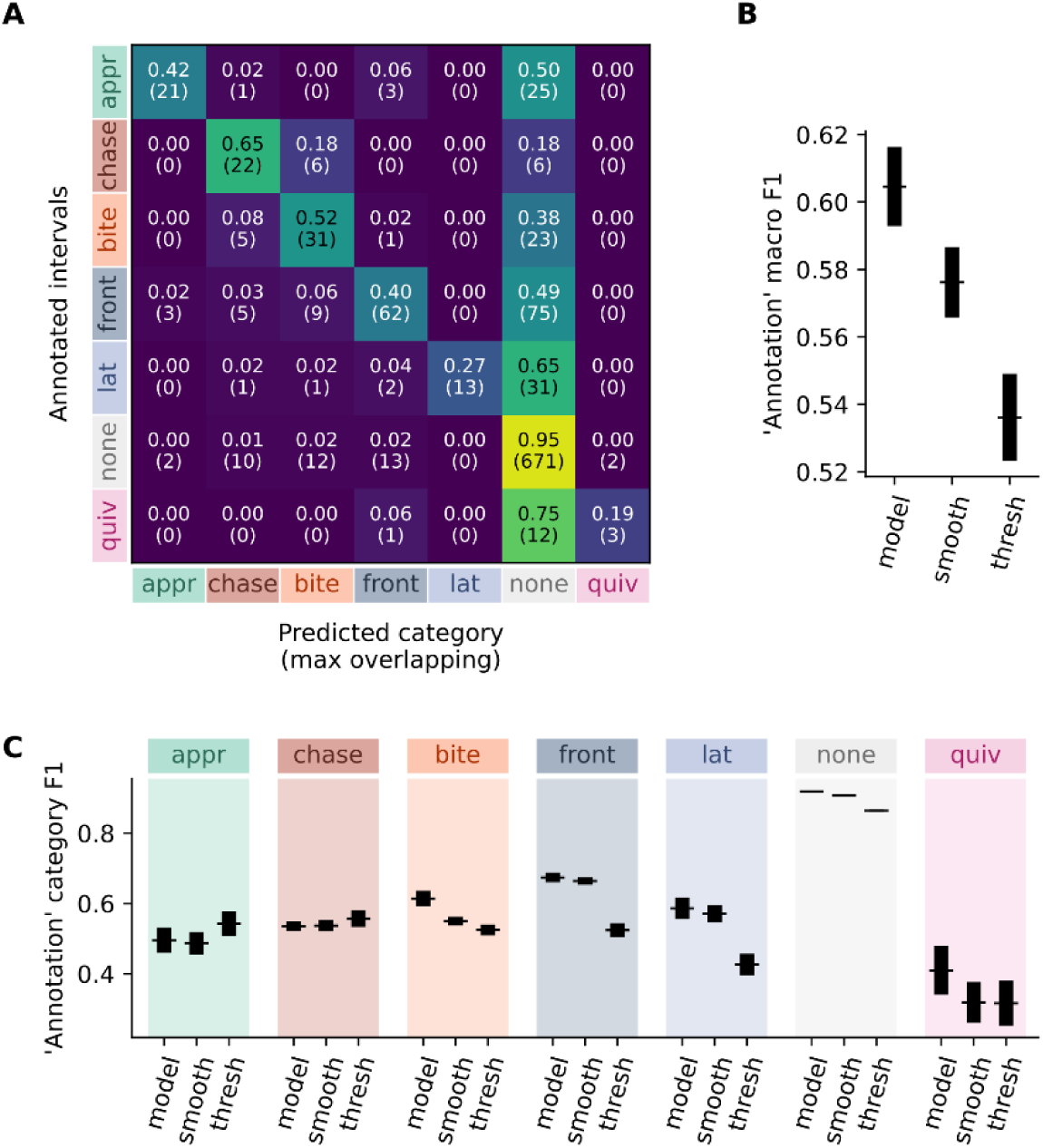
Additional social cichlids test dataset validation results based on annotated intervals. The predicted category that corresponds to a behavioral annotation is selected as the category of the predicted interval with longest overlap. All values represent means of 20 pipeline runs with different random states, standard deviations are shown as error bars if applicable. **A:** Confusion matrix depicting recall of annotations. Note that the ‘none’ column represents false negatives (i.e., missed predictions). Proportional values are normalized across rows, absolute counts are shown below in parentheses. **B and C:** Macro F1 and category F1 scores calculated on annotated intervals for three (post-)processing steps – raw model outputs (‘model’), after probability smoothing (‘smooth’) and after thresholding (‘thresh’).

**SI Figure 3:**
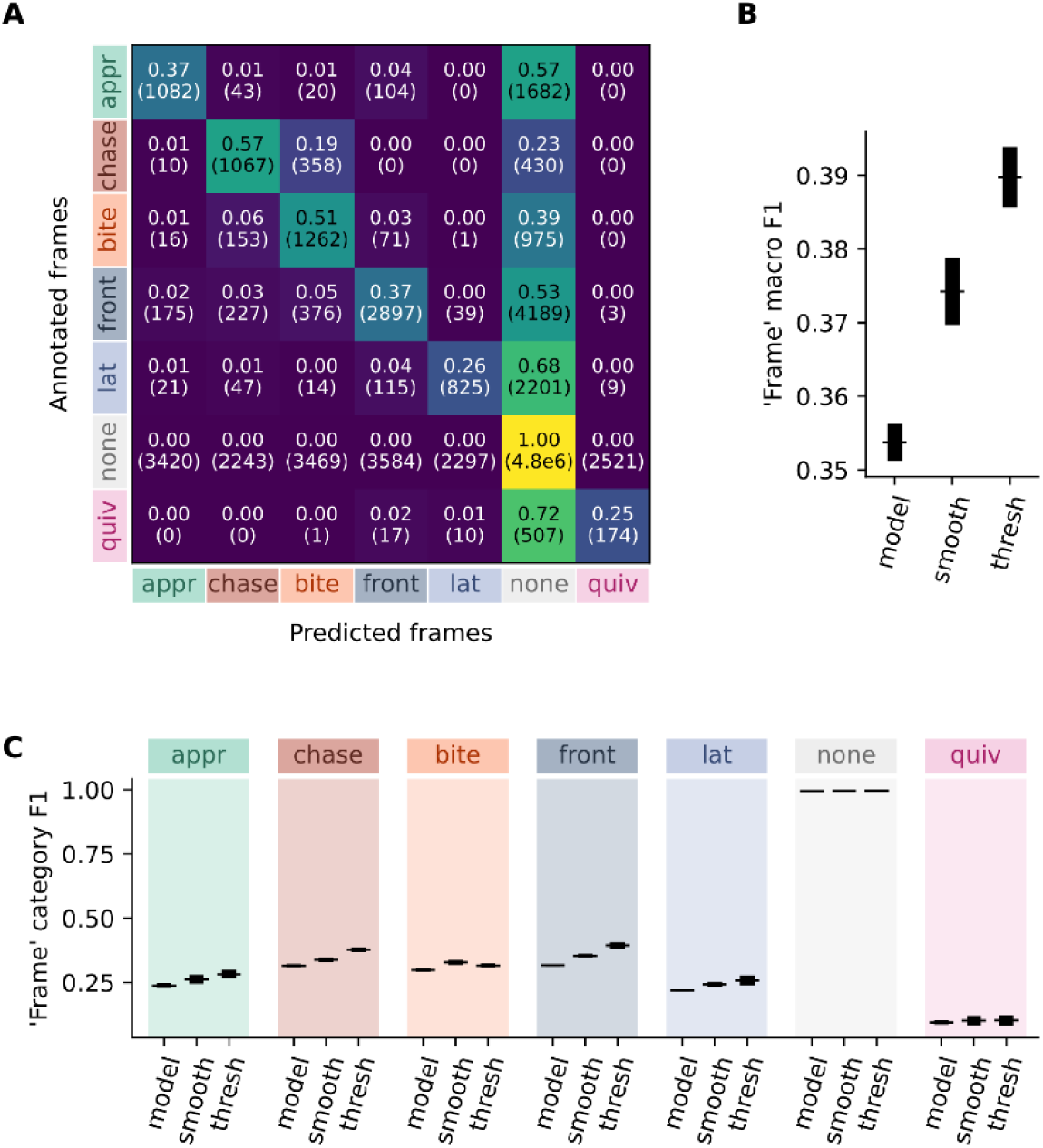
Additional social cichlids test dataset validation results based on video frames. All values represent means of 20 pipeline runs with different random states, standard deviations are shown as error bars if applicable. **A:** Confusion matrix across all video frames of the test dataset. Proportional values are normalized across rows, absolute counts are shown below in parentheses. Note the disproportionate number of frames that belong to the behavioral background category ‘none’. **B and C:** Macro F1 and category F1 scores calculated on video frames for three (post-)processing steps – raw model outputs (‘model’), after probability smoothing (‘smooth’) and after thresholding (‘thresh’).

**SI Figure 4:**
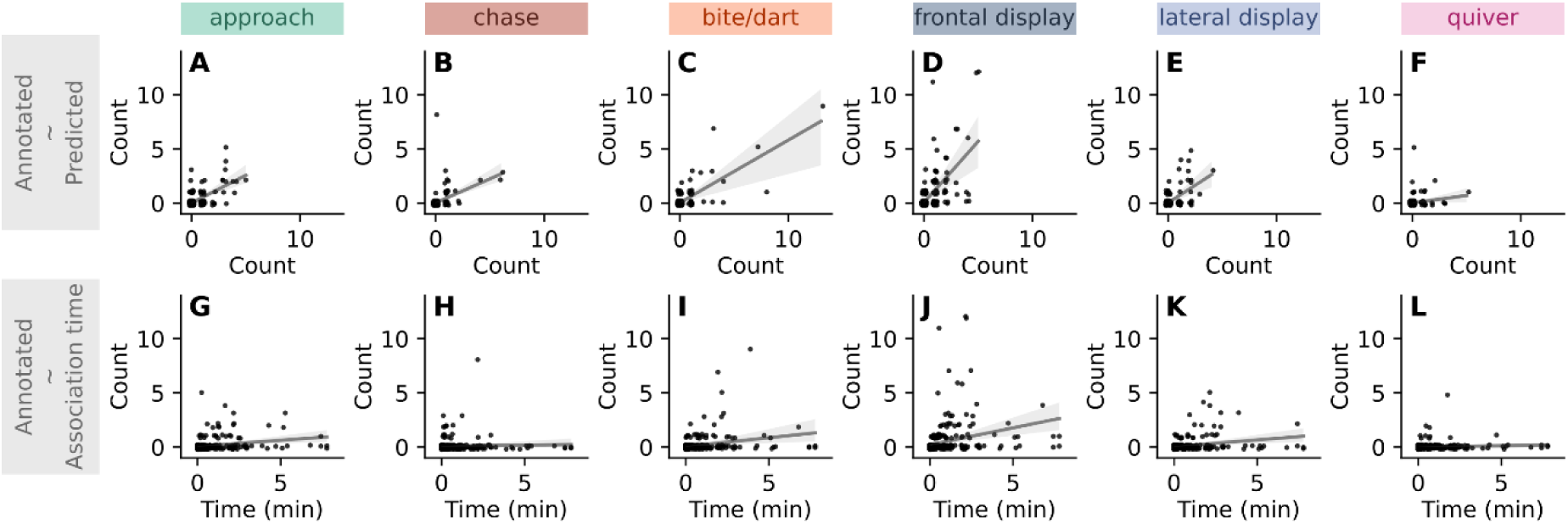
Visualization of correlations between ground-truth, annotated behavioral interaction counts and two potential behavioral proxies: **A – F:** Correlations with predicted counts, split by behavioral foreground categories. **G – L:** Correlations with association time, the cumulative duration that two individuals spent within a distance of three average body lengths. The correlation strengths of all correlations with predicted counts are significantly higher than the corresponding correlation strengths with association time (William’s correlation tests with dependent correlations that share one variable; P < 0.001 in all cases). For statistical estimates and a sensitivity analysis with other association distance thresholds (1 and 5 body lengths), see SI Table 3.

**SI Figure 5:**
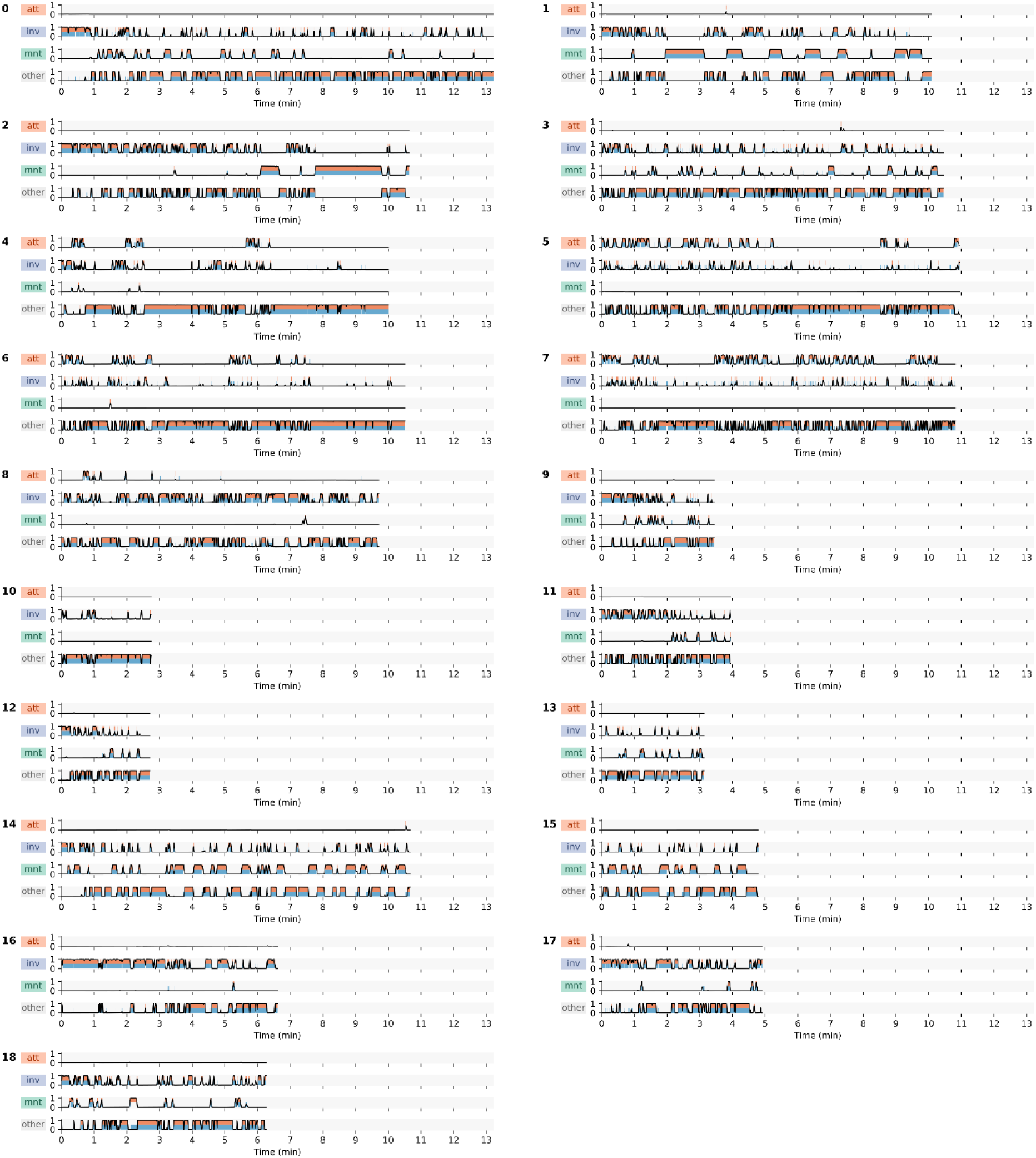
Classification results of all 19 resident-intruder sequences in the CALMS21 test dataset after post-processing (i.e., category-specific output smoothing and thresholding). Upper bars (orange) show predicted intervals, lower bars (blue) ground-truth annotations. Lines represent model outputs – classification probabilities for each category – after smoothing. All sequences are visually aligned with the longest sequence ‘0’. Note the skewed distribution of annotated ‘attack’ intervals: Only five of 19 sequences have annotations for this behavioral category, but in those at a relatively high frequency.

### Supplementary Videos

**SI Video 1**: Example usage of the interactive validation tool. See Figure 4 for more information. The video and subtitles are also available in the online documentation at vassi.readthedocs.io. Note that Figure 1 as shown in the video depicts classification results from a preliminary classifier that was trained with a slightly different subsampling strategy.

